# Forces experienced by instrumented animals depend on lifestyle

**DOI:** 10.1101/2020.08.20.258756

**Authors:** Rory P Wilson, Kayleigh A Rose, Richard Gunner, Mark Holton, Nikki J Marks, Nigel C Bennett, Stephen H. Bell, Joshua P Twining, Jamie Hesketh, Carlos M. Duarte, Neil Bezodis, D. Michael Scantlebury

## Abstract

Animal-attached devices have transformed our understanding of vertebrate ecology. However, to be acceptable, researchers must minimize tag-related harm. The long-standing recommendation that tag masses should not exceed 3% of the animal’s body mass ignores tag forces generated by movement. We used collar-attached accelerometers on four free-ranging carnivores, spanning two orders of magnitude in mass, to reveal that during movement, forces exerted by ‘3%’ tags were generally equivalent to 4-19% of the animals’ masses, with a record of 54% in a hunting cheetah. Controlled studies on domestic dogs revealed how the tag forces are dictated by animal gait and speed but appear largely invariant of body mass. This fundamentally changes how acceptable tag mass limits should be determined, requiring cognizance of animal athleticism.

**One Sentence Summary:** There can be no universal rule for collar-tag masses as a percentage of carrier mass since tag forces depend on lifestyle.

## Main Text

The use of animal-attached devices is transforming our understanding of wild animal ecology and behavior (*1, 2*). Indeed, smart tags have been used across scales to measure everything from the extraordinary details of high performance hunts in cheetahs (*3*), to vast cross-taxon comparisons of animal behavior and space-use over whole oceans ((e.g. *1, 4*)). A critical proviso is, however, that such devices do not change the behavior of their carriers, for both animal welfare issues as well as for scientific rigor (*5*). Defining acceptable device loads for animals is critical because even diminishingly small tags can cause detriment. For example, Saraux et al (*6*) showed that the addition of flipper rings to penguins can affect their populations, presumed to be due to the tags increasing the drag force in these fast-swimming birds. Performance is relevant in this case because drag-dependent energy expenditure to swim increases with the cube of the speed (*7*).

Although consideration of the physics of drag has been shown to be a powerful framework with which to understand tag detriment in aquatic animals (e.g.(*8*) (*9*)), drag is negligible in terrestrial (though not aerial) systems even though tag detriment in terrestrial animals has been widely reported and is multi-facetted (*10*). For example, cited issues range from minor behavioral changes (*11*) through skin-, subcutaneous- and muscle damage with ulceration *12, 13*) to reduced movement speed (*14*) and dramatically increased mortality (*15*). As with drag, we advocate that a force-based framework is necessary to help understand such detriment. Indeed, force is implicit in ethics-based recommendations for acceptable tag loads because, for example, a central tenet is that animal tags should never exceed 3% or 5% of the animal-carrier body mass (*16*). Importantly though, we could find no reason for advocating this value, which thus remains arbitrary. Most likely, it was based on the widely accepted 5% significance value used in biological statistics. Implicit in this limit is that consequences, most particularly the physical forces experienced by animals due to tags, are similarly limited. This cannot be true because Newton showed that mass, force and acceleration are linked *via* F = ma, so animal performance, specifically their acceleration, will affect the tag forces applied to them. Tag forces on their animal carrier can therefore be accessed by measuring acceleration experienced by the tag as the animal moves. We note here though, that this necessitates gathering on-animal data because simple consideration of acceleration from rigid-non-living bodies is inappropriate for living systems composed of multiple interacting segments (*17*).

Here, we examine the forces exerted by collar-mounted tags on moving animals; lions *Panthera leo*, European badgers *Meles meles*, pine martens *Martes martes* and a cheetah *Acinonyx jubatus* (with body masses roughly spanning 2–200 kg) equipped with accelerometers undertaking their normal activities in the wild. We also equipped twelve domestic dogs *Canis familiaris* (2-45 kg) with the same tags but with masses of up to 3% of the dog body mass as they travelled at defined speeds. In both instances, we examined how the acceleration changed with animal motion and then calculated the forces experienced by the animals due to the tags. In all instances of movement, the tag forces rose, increasing with animal speed and athleticism, escalating in some instances to be much greater than 20% of the carrier’s normal body mass. This fundamentally changes how we should view critical limits of tag masses.

Accelerometer data, summarized as the vectorial sum of the three orthogonal axes, from the equipped animals during movement showed tri-modal distributions except for the pine martens which were mono-modal. Following (*18*) we considered that these most likely corresponded to walking, trotting and bounding (cf. our direct observations of the domestic dogs below); these were further exemplified by variation in the amplitude and periods of peaks in this acceleration metric (Fig. 1). Cumulative frequencies of all acceleration values showed increasing acceleration from walking through trotting to bounding and typically had a roughly logarithmic-type curve for all gaits and animals (Fig. 1). The percentage time during which the tags carried by the carnivores had acceleration exceeding 1 *g* varied between a mean minimum of 31% for walking badgers to 88% for bounding cheetahs (Table S1). Furthermore, while species differences in acceleration distributions were not readily apparent for their walking gaits, the percentage time during which acceleration was in excess of 1 *g* was greatest during bounding, with cheetahs showing the highest values in this category (Fig. 1). Mean peak accelerations per stride across species varied between 1.37 *g* (SD 0.05) and 6.25 *g* (SD 0.79) for walking and bounding cheetahs, respectively (Table S2). The maximum recorded value was 18.1 *g* in a cheetah assumed to be chasing prey.

**Fig. 1.**
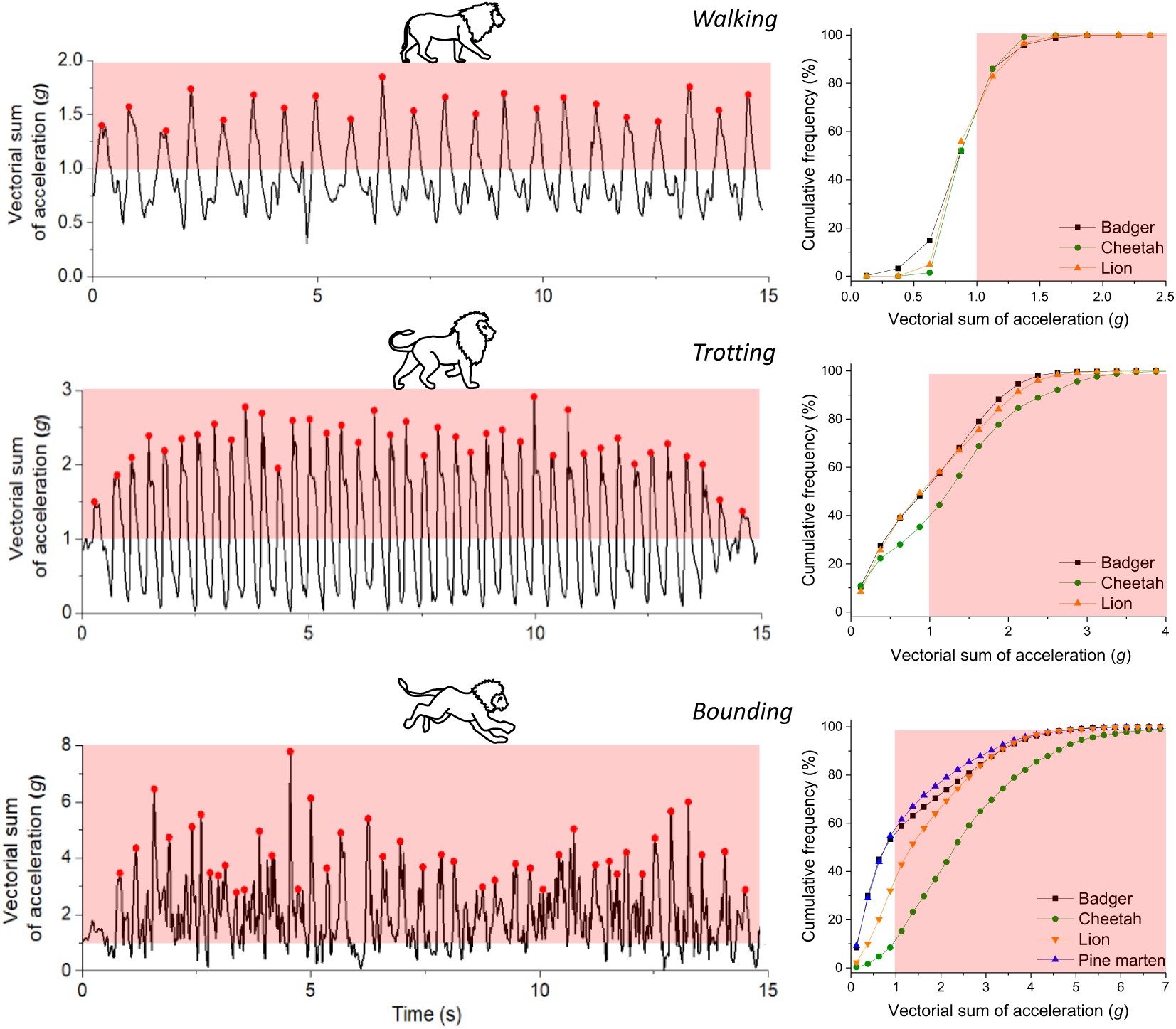
Acceleration signatures vary according to gait and lifestyle. Left-hand panels; Acceleration signatures recorded by collar-mounted tags on a lion according to activity. The red areas show when the acceleration exceeded that of gravity (note the changing scales with gait). Right-hand panels; Cumulative frequency of all acceleration values for four free-living carnivores according to gait.

Across species, gait was the main factor dictating peak acceleration (Fig. 2) and there were no significant effects of body mass, nor period between peaks (linear mixed-effects model: log period: *F*_1, 210_= 0.01, *P*=0.908; gait: *F*_2, 208_=1083.07, *P*<0.0001; body mass: *F*_1, 19_= 3.00; *P*=0.100, Table S3). The period between acceleration peaks was greater for larger species during slower gaits, but not for bounding (a linear mixed-effects model demonstrated a significant interaction effect between body mass and gait: *F*_2, 209_= 3.00, *P*<0.0001, Table S3).

**Fig. 2.**
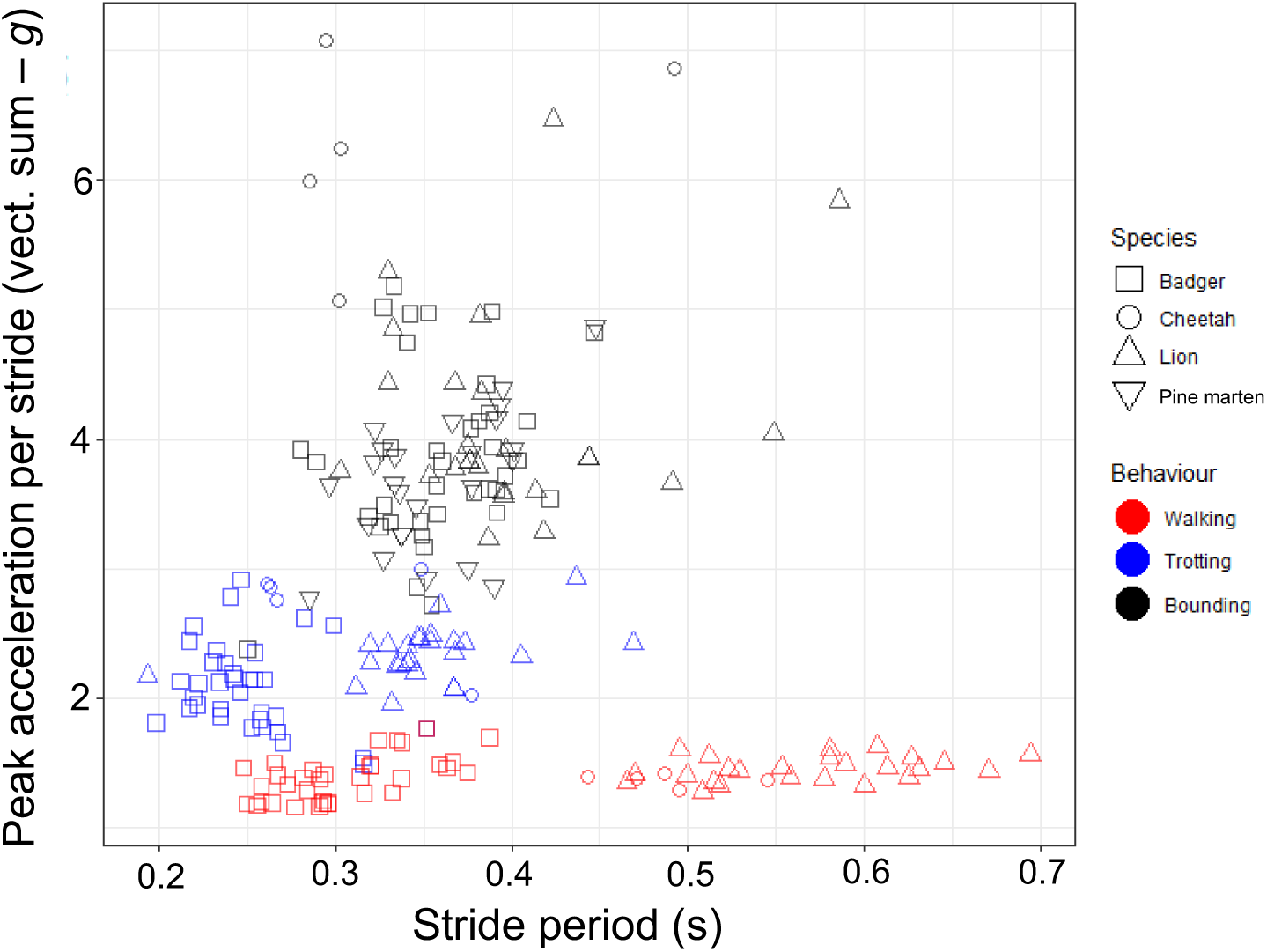
Body mass and stride period do not dictate peak tag acceleration. Peak amplitudes of (the vectorial sum of) accelerations versus stride periods for four free-living carnivores (see symbols) travelling using different assumed gaits (colors). Each individual point shows a mean from a duration of activity >5 s from a single individual. See also Fig. S1 for similar data from domestic dogs.

Acceleration metrics changed even within particular gaits though, as exemplified by predators chasing prey. For example, overall, prey chases exhibited by lions showed mean peak accelerations increasing from 1 *g* to mid-chase peaks of *ca*. 5 *g* before decreasing again (Fig. 3). Assuming that these animals were carrying tags that amounted to 3% of their mass, this would result in forces amounting to up and beyond 15% of their normal body mass (e.g. Fig. 3). The maximum observed was 54% in the cheetah.

**Fig. 3.**
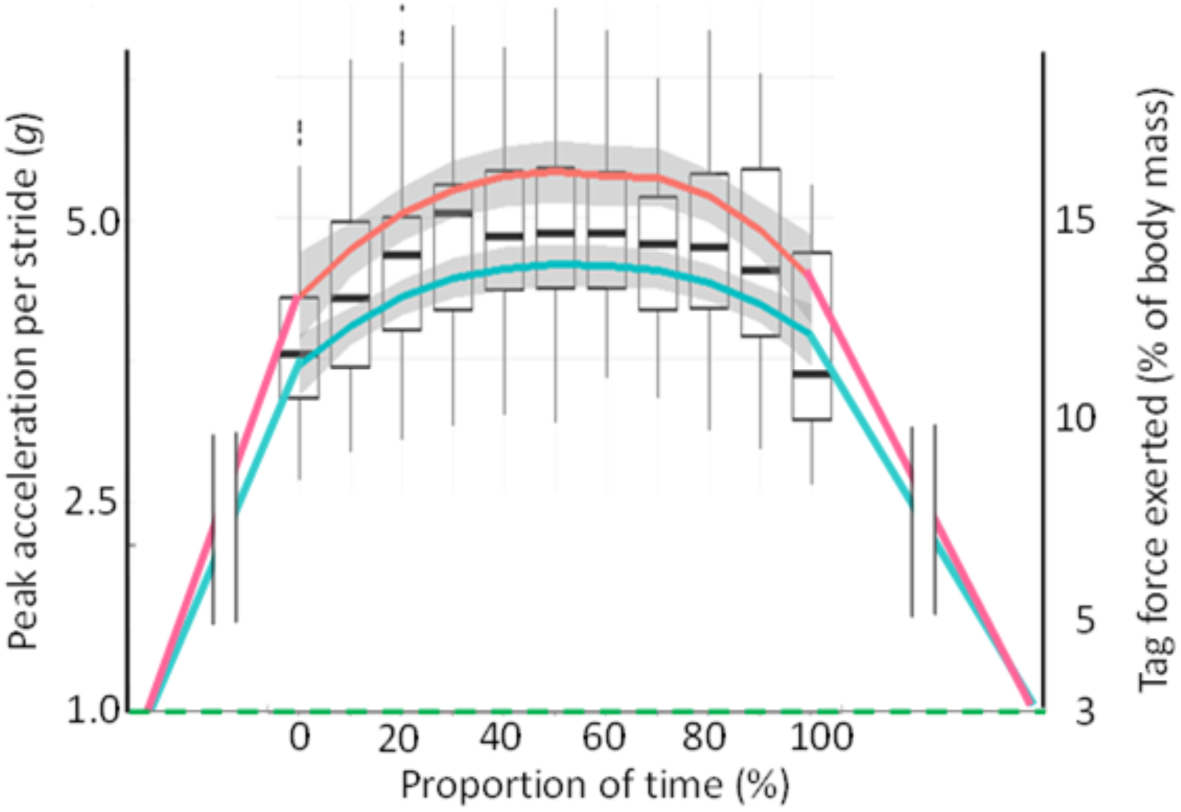
Hunting lions experience maximum tag forces mid-chase. Box whisker plot [bold horizontal bars show means and boxes inter-quartile ranges] of the changing forces (the vectorial sum of the acceleration peaks per bound [cf. Fig. 1] and as a percentage of animal body mass assuming the tag constitutes 3% during non-movement) exerted by animal-attached tags on lions chasing prey as a function of the progression of the chase. Red and blue lines show grand means for 5 females and 5 males, respectively. The maximum acceleration was > 12.5 g, equating to a 3% tag exerting a force equivalent to 37.5% of the animal’s normal weight.

In dogs, stride peak accelerations increased linearly with travel speed (Fig. 4) but at greater rates with increasing relative tag mass above tag masses equivalent to 1% of carrier body mass (there was a significant interaction effect between travel speed and tag % body mass: *F*_3, 500_ = 4.44, *P*=0.004, *R*^2^ = 0.74 Table S4). There was also a significant interaction effect between gait and tag mass as a percentage of carrier body mass (*F*_6, 498_ = 4.33, *P*=0.0002, *R*^2^ = 0.74, Table S4). Peak accelerations ranged from 4-18 *g* (fig S1-2) during bounding with collar tags equivalent to 3 % of the carrier body mass. Movement of the tag relative to the collar and body (flapping/ swinging) was exacerbated under this condition and, as a consequence, the force exerted by the tags ranged from 20-50% of normal carrier body mass.

**Fig. 4.**
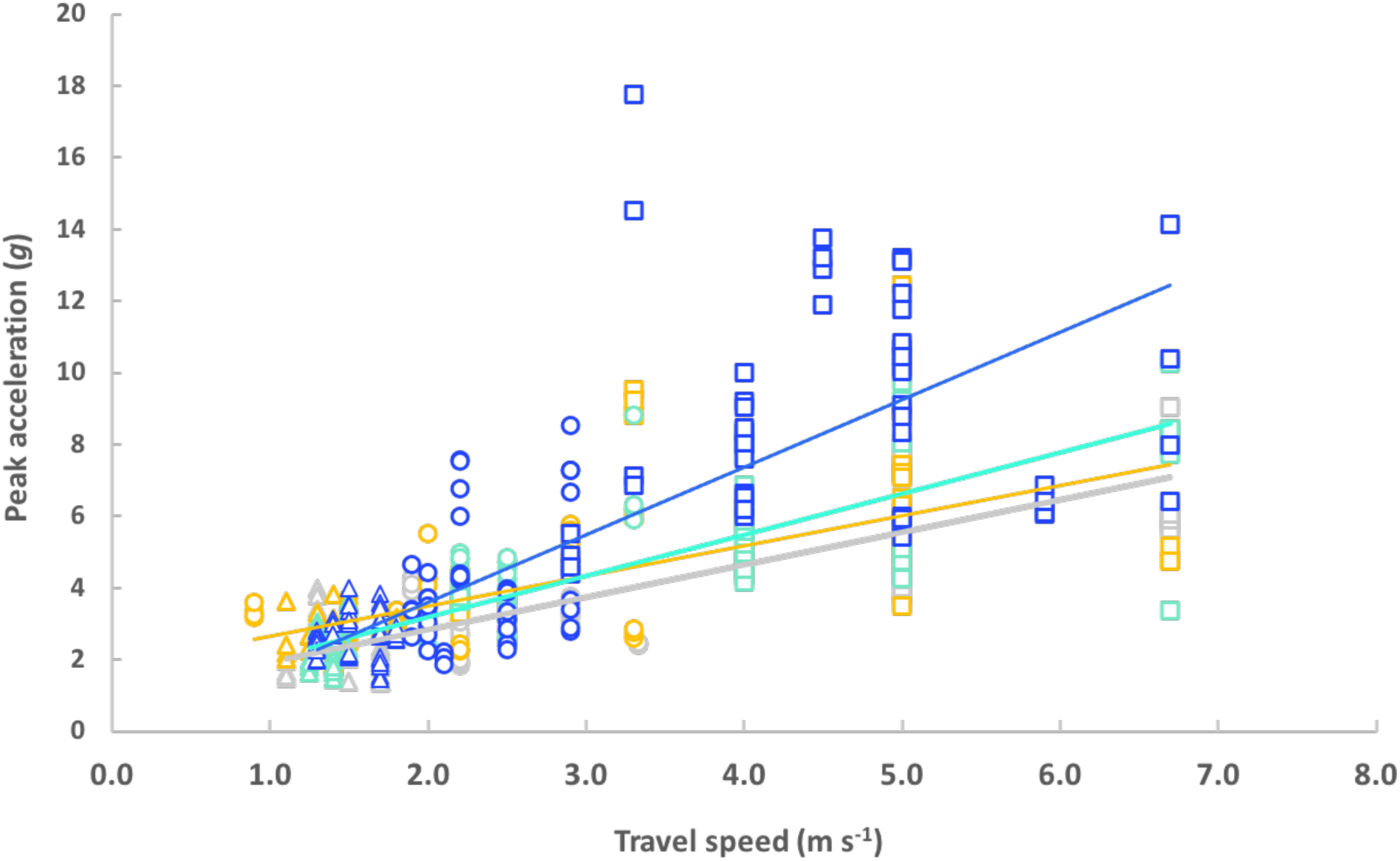
Travel speed and tag mass influence tag peak acceleration. The relationship between peak accelerations and travel speed for 12 individual dogs (masses 2-45 kg), colored according to the percentage mass of the tag relative to the carrier; 0 (grey, *y = 0.90x + 1.04*), 1 (yellow, *y = 0.84x +1.80*), 2 (light blue, *y = 1.15x + 0.88*), 3 (dark blue, y = 1.88x -0.16). Data points are the means of the four greatest peaks in acceleration per 20 m trial per gait and dog. Data for walking, trotting and bounding gaits are represented as triangles, circles and squares, respectively. Coefficients for the best-fit lines are taken from the final model outputs (Table S4).

Stride peak accelerations were largely invariant with body mass (*F*_1,10_ = 3.51, *P*=0.09, Table S4) across dog breeds for any given gait (Fig. S2). Consequently, the peak forces exerted by the tags were directly proportional to tag mass and body mass. Accordingly, relative forces (force as a percentage of normal carrier body mass) were independent of carrier body mass (Table 4, Fig. S3).

## Discussion

A vehicle accelerating in a straight-line only experiences acceleration in the longitudinal axis. In contrast, the multiple limb-propelled motion of an animal with a flexible body produces complex trunk accelerations owing to the changing limb accelerations (*17*), synergistic effects of limb joint kinematics and limb muscle groups that contribute to the reduction and transfer of mechanical energy effecting shock absorption (*19*), and the mechanical work conducted within each stride (*20*). Ultimately, the magnitude of trunk accelerations depends on the combined variation in acceleration of the limbs, and the masses of those limbs (cf. *21*). Thus, animals engaging in high performance activities are expected to produce high body accelerations, and have physiological and anatomical adaptations to enhance performance, such as fast twitch muscles (*22*), and energy stored within tendons (*23*) for explosive release, which will increase this. Through all these complexities, tags mounted on the trunk of an animal result in extra forces being imposed that scale linearly with the acceleration of the tag and its mass. Consideration of animal lifestyle then, can already inform prospective tag users of the likely scale-up of the tag forces beyond the 1 *g* normally considered for tag detriment. Consequently, the 3-5% mass limits for slow-moving animals, such as sloths (Bradipodidae), seem most appropriate, while they may not be for pursuit predators, such as wild dogs (*Lycaon pictus*), regularly jumping animals like kangaroos (Macropodidae) or rutting ungulates (Ungulata). Beyond that, in our small sample of carnivores at least, which nonetheless covers about two orders of magnitude in mass, it seems that peak acceleration associated with gait varies little with mass, although larger animals have longer stride periods (Fig. 2 – cf. (*24*)). If these animals were to carry tags constituting 3% of their normal body mass, mean peak forces imposed by the tags would constitute *ca*. 4.5%, 6% and 12% of this body mass for walking, trotting and bounding gaits at frequencies of between 1.6 and 4 times per second (for walking lions and trotting badgers, respectively - Fig. 2, Table S2). Importantly, tag attachment is relevant in translating the acceleration experienced by the animal’s trunk into tag-dependent forces acting on the animal, with collars predicted to be particularly problematic. A tag that couples tightly with its carrier’s trunk, such as one attached with tape to a bird (*25*) or glue to a marine mammal (*26*), experiences acceleration that closely matches that of its substrate, so it exerts forces at a site where most of the animal’s mass lies. In contrast, a device on a looser-fitting collar of a moving tetrapod not only exerts forces on the (less massive) head and neck areas, rather than the animal trunk, but the tag also oscillates between essentially two states: One is analogous to ‘freefall’, which occurs between pulses of animal trunk acceleration in the stride cycle which project the collar in a particular direction owing to its inertia and lack of a tight couple with the neck. The collar is therefore subject to peaks in acceleration when it interacts with the animal’s neck, causing greater collar acceleration than would be the case if it were tightly attached to the animal’s body (cf. peaks in Fig. 1). This explains why Dickinson et al. (*27*) reported that acceleration signatures from collar-mounted tags deployed on (speed-controlled) goats *Capra aegagrus* became increasingly variable with increasing collar looseness, and is analogous to the concerns related to injuries sustained by people in vehicles depending on seatbelt tightness (*28*). Partial answers to minimizing such problems may involve having padded collars that should reduce acceleration peaks, making sure that the tags themselves project minimally beyond the outer surface of the collar and having wider collars to reduce the pressure.

Having identified how animal movement changes the 3-5% tag rule, it is more problematic to understand how the identified forces translate into detriment. A prime effect is that higher forces and smaller contact areas will lead to higher pressure at the tag-animal interface because pressure = force/area. This can affect anything from fur/feather wear (*29*) to changing the underlying tissue (*30*) and, as would be predicted, is notably prominent in highly active species wearing thin collars (e.g. Howler monkeys *Alouatta palliatai*, where 31% of animals wearing ball-chain radio-collars constituting just 1.2% of their mass sustain severe damage extending into the subcutaneous neck tissue and muscle (*12*)). But pressure-dependent detriment will also depend on the proportion and length of time to which an animal is exposed to excessive forces, with animals that spend large proportions of their time travelling, such as wild dogs, being particularly susceptible (*31*).

Perhaps more esoteric though, is the extent to which the inertia of a variable force-exerting tag ‘distracts’ its wearer, aside from the physical issues of load-bearing by animals. The tag mass as a percentage of carrier mass did not affect the gait-specific speeds selected by the domestic dogs in this study. However, it remains to be seen the extent to which a typical 30 kg cheetah wearing a collar that is 3% of its body mass, and therefore experiencing an additional force equivalent to up to 16 kg during every bound of a prey pursuit, might have its hunting capacity compromised. We note that the survival of such animals is believed to be especially sensitive to the proportion of successful hunts (cf.(*32*)) which calls for critical evaluation of performance between tag-wearing and unequipped animals, or animals equipped with tags of different masses (cf.(*33*)). In the meantime, we suggest two approaches to assessing likely tag force effects for planned tag deployments on animals. Firstly, simple consideration of animal lifestyle, where possible backed up by (tri-axial) acceleration metrics, will go some way to getting a more realistic assessment of potential for detriment. Secondly, we suggest that, where acceleration data are available from wild animals, consideration of a frequency histogram of the vector sum of the tag acceleration (Fig. 5) will help indicate the extent and intensity of the force interactions between the animal and its tag.

**Fig. 5.**
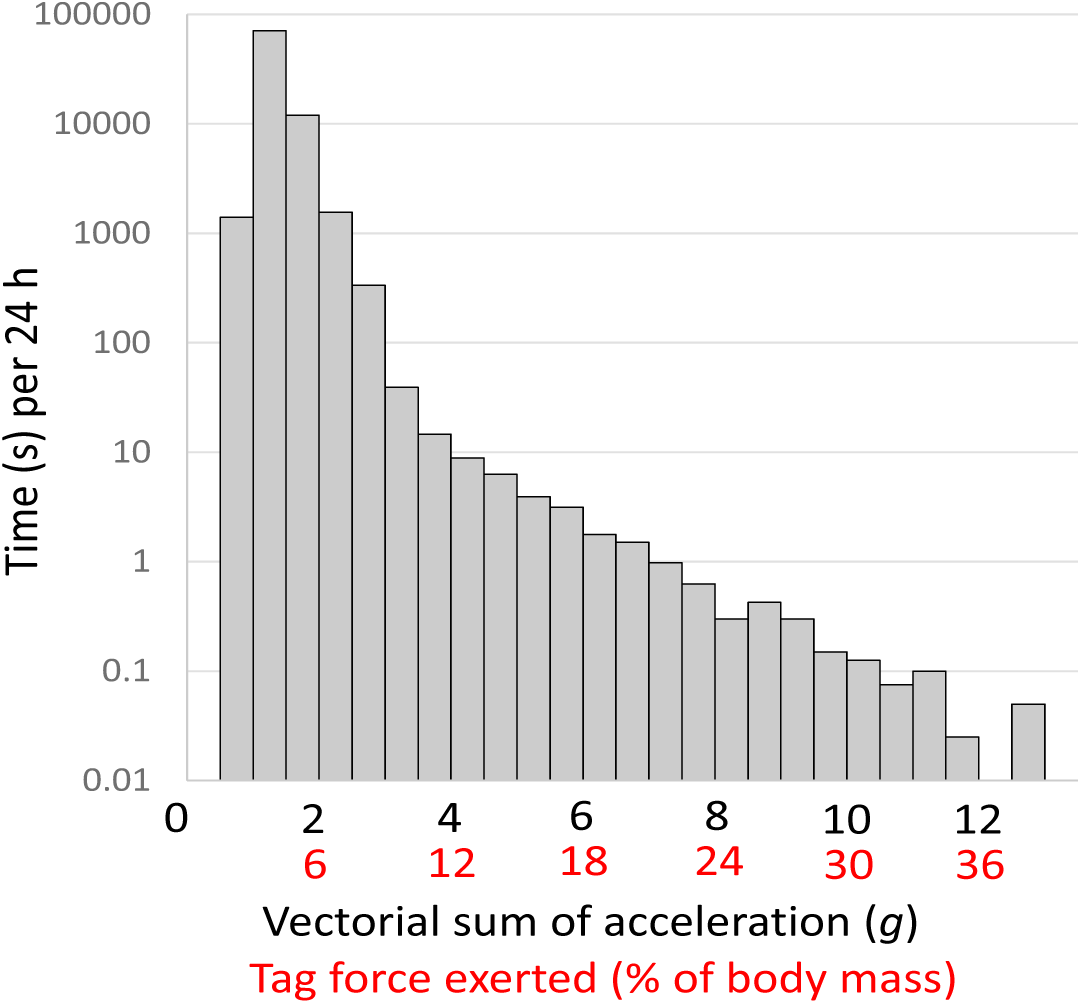
Tag forces on their animal carriers can be expressed in a time histogram. Time (expressed as the log of the time) that a badger-attached tag spent subject to different accelerations over 24 h. The diagram also converts the acceleration values into forces imposed on the badger as a percentage of its normal body mass assuming the tag constitutes 3% during non-movement (axis in red). Although higher g-forces are a diminishingly small proportion of the badger’s time, their detrimental effect may not be a linear function of their values.

Small contributions in time to high forces should not be ignored simply due to their fractional components. Such issues can likely make the difference between a predator catching its prey or not, or worse, if the prey is wearing the tag, may result in it being caught.

Thus, while the contributions of animal-attached tag technology to advance scientific understanding are unquestionable, simple consideration of acceleration signatures of animal movements can go a long way to helping us understand how the effects of such tags on the behavior of their carriers vary with activity. Most importantly, these considerations should enable us to develop systems that minimize such effects, to the benefit of both animals and the science that their studies underpin.

## Acknowledgments

We gratefully acknowledge the access provided by the National Trust and Forest Service NI. We are also grateful to the RSPCA’s Llys Nini Wildlife Centre in Penllergaer, Wales, and to Judy Corbett and Peter Welford at Gwydir Castle in Llanrwst, Wales, for allowing us to work on their property with their dogs. We thank Derek van Heerden and the staff at Harnas Wildlife Foundation in Namibia for their kindness and supporting our work. We thank SANParks and the Department of Wildlife and National Parks, Botswana for allowing our research in the Kgalagadi Transfrontier Park (Permit Number SCAM 1550). We are grateful to Angela Bruns, Sam Ferreira and Danny Govender and Pauli Viljoun for facilitating the research and to the many field staff and volunteers that conducted the fieldwork including Wayne Oppel, Corera Links, Martin van Rooyen and Mads Frost Bertelsen. We are grateful to Fraser Menzies and all the field staff at the Department of Agriculture, Environment and Rural Affairs, DARD. We are also grateful to the Vincent wildlife trust for supporting this research.

## Author contributions

RPW conceived the work and wrote the manuscript with KAR and DMS. RG, SHB, NM, JPT, J.H and DMS collected data, which was analyzed by RPW, KAR, RG and DMS using software developed by MH. All authors provided valuable input to the writing and primary analysis.;

## Data and materials availability

All data will be made available on Figshare following acceptance for publication.

## Supplementary Materials

Materials and Methods

Figures S1-S3

Tables S1-S5

Reference (*33*)

