## Supplementary methods for "Forces experienced by instrumented animals depend on lifestyle"

Materials and Methods

Tag deployments on wild species

We selected 4 species of free-living carnivores, exemplifying about 2 orders of magnitude of mass; 10 lions (mean mass *ca.* 180 kg), 1 cheetah (mass *ca.* 40 kg), 10 badgers (mean mass *ca.* 8 kg) and 5 pine martens (mean mass 1.9 kg), and fitted them with collar-mounted tri-axial accelerometers (‘Daily Diaries - Wildbyte Technologies [<http://www.wildbytetechnologies.com/>]; measurement range 0-32 *g*, recording frequency 40 Hz). Due to the weighting of the loggers, and more particularly their associated batteries, the units and sensors were normally positioned on, or close to, the underside of the collar although during movement the collars could rotate. After being equipped, the animals roamed freely, behaving normally, for periods ranging between 1 and 21 days before the devices were recovered.

Badgers were live-trapped in various locations in Northern Ireland in custom-built cages and anaesthetised with a mixture of ketamine, medetomidine and butorphanol IM with recumbency achieved 5-10 minutes post injection. Loggers were attached to an adjustable nylon clip-on dog collar (Ancol Pet Products Limited, Walsall; circumference 20-30 cm) with a layer of waterproof self-amalgamating tape (ultratape; Bruce Douglas Marketing, Dundee, UK), which was additionally fastened with three cable ties and then covered with ‘tesa’ tape (No. 4651; tesa AG, Hamburg, Germany). Badgers were recaptured approximately 10 days later, and the collars removed.

The cheetah work took place at Harnas Wildlife Foundation in Namibia. A hand-reared individual was used, so a collar could be placed around its neck without anaesthesia. Once released, the individual was free to move and hunt at will. Data were downloaded each day. We used a commercially available nylon collar (EzyDog Medium dog collar, circumference 29-40cm, Pets at home, Handforth, England) to which we attached the logger with three cable ties and tesa’ tape as described above.

Lions were captured from the Kgalagadi transfrontier park in South Africa and treated according to SANParks operational procedures as detailed in SANPark’s ‘Standard Operating Procedures for the Capture, Transportation and Maintenance in Holding Facilities of Wildlife’ (Reference: 17/Pr-CSD/SOP capture, transport, holding facilities (04-17) v2) and SANParks SOP ‘Fitment of tracking devices and marking of fauna in South African National Parks’ (Reference: 17/Pr-CSD/pro/tracking + marking (12/16) v1). A pride of lions was identified during the course of the day through field rangers’ observations and tourist sightings. At nightfall a bait station was set up nearby the pride, consisting of an antelope carcass secured to a tree. This carcass was freshly acquired during the day by authorized ranger staff. The intestines were removed via a cut through the abdominal midline and intestinal contents used to lay a scent trail towards the bait. The darting vehicle was positioned at 20 m distance from the bait station and lion alerted and lured towards the bait by broadcasting pre-recorded animal distress calls via a long-range loudspeaker system (Truck Pro, Foxpro Inc). Once lions settled to feed, the targeted individuals (all adults) were identified and their weight estimated. Subsequently the animals were darted with a DAN-INJECT CO2 injection rifle (Model JM.SP.25, DAN-INJECT). A drug combination of zolazepam/tiletamine (Zoletil®Virbac) and medetomidine hydrochloride (Medetomidine Compound, Kyron Laboratories,) at an average 1.2 mg/kg and 0.05 mg/kg respectively as adapted from recommendations in SANPark’s ‘Standard Operating Procedures for the Capture, Transportation and Maintenance in Holding Facilities of Wildlife’ and Kock *et al. ^33^* was used. The darted individuals were followed visually with the help of spotlights. Once animals became recumbent they were recovered, secured with blindfold and foot shackled and translocated to a nearby processing station. They were monitored for temperature, respiration and heart rate. Collars were fitted around the neck in accordance with SANParks SOP ‘Fitment of tracking devices and marking of fauna in South African National Parks’, allowing for three fingers space and ensuring collar size did not exceed the maximum head circumference. Where prolonged anaesthesia was required, animals were given an additional increments of ketamine (Ketamine Powder Compound, Kyron) intramuscularly at a total average of 1.44mg/kg. Dart sites were treated systemically via subcutaneous injection at the recommended dose with Ceftiofur (Excede®, Zoetis,, 1ml per 30 kg) and meloxicam (Metacam®, Boehringer, Randburg, 0.2mg per kg) respectively. Once processed, animals were relocated close to the pride or capture location and the medetomidine component of anaesthesia reversed intramuscularly with a mix of atipamezole (Antisedan®, Zoetis) at 2.5 times the amount of medetomidine and yohimbine (Yohimbine Compound, Kyron) at 6.25 mg per kg body weight. Animals were monitored until ambulatory. All procedures followed recommendations of SANPark’s ‘Standard Operating Procedures for the Capture, Transportation and Maintenance in Holding Facilities of Wildlife’.

Pine martens (males) were live-trapped and equipped with tags in Northern Ireland between November 2018 - March 2019, and August - November 2019. Devices were attached to the collar using self-amalgamating tape. The complete system amounted toa collar weight of approximately 45g (*ca.* 2.3% of bodyweight). Deployments lengths ranged from 5 – 14 days

Trials with domestic dogs

Twelve domestic dogs (*Canis lupus domesticus*) of seven different breed combinations and three main body types (small, racers and northern breeds), ranging 2-45 kg in body mass (Table S5), were volunteered by their owners and the RSPCA’s Llys Nini Wildlife Centre (Penllergaer, Wales) to take part in this study (Table S5). Dog body masses were provided by owners and body length, forelimb length and hindlimb length were measured to the nearest cm. Two leather dog collars (short and long) of the same width were used to cover the range in dog neck size. Combinations of pre-prepared lead plates (up to 10 cm in length) and varying in mass (25, 35, 45, 50, 100, 150 and 175 g) were fashioned into collar loads equivalent to 1, 2 and 3% of each carrier dog’s body mass. The loads were stacked and attached securely to the ventral collar along their full-length using Tesa® tape. A tri-axial accelerometer (Daily diary, measurement range 0-32 *g*, recording frequency 80 Hz, Wildbyte Technologies) and its supporting battery (3.2 V lithium ion) were taped securely to the load. The tag and battery combined weighed 11.9 g and, in the absence of any additional load, were considered negligible in mass and used as a control (0 % carrier body mass). All trials were approved by the Swansea University Animal Welfare Ethical Review Body (ethical approval number IP-1617-21D).

All trials were filmed and each dog was encouraged to walk, trot and bound along a 25 m stretch of level, short-cut grass wearing collar tags equivalent to 0, 1, 2 and 3% of their body mass (twelve gait and tag mass combinations) and trial order was randomised. A stopwatch was used to record the time taken (to the nearest s) for a dog to travel 20 m between appropriately spaced posts in order to calculate an average speed of travel (m s^-1^).

Data processing

In both the cases of the free-living carnivores and domestic dogs, the 3 channels of raw acceleration data were converted to a single channel by calculating the vectorial sum of the acceleration following Vect sum = √(a_x_^2^+a_y_^2^+a_z_^2^), where a is the instantaneous acceleration and the subscripts denote the different (orthogonally placed) acceleration axes. The specifics of the surge, heave and sway accelerations were not considered separately due to some collar roll. We selected 4 peak accelerations from the gait waveforms to examine as a function of speed, gait, body mass and tag mass as a percentage of carrier body mass in the dogs (SI 2). We standardized the use of four peaks because at the highest speeds some dogs only had four full waveforms during the test stretch. Gait was assessed visually in the dogs as a walk, trot or gallop. The forces exerted by the tags on their animal carriers were calculated using F = ma, where m is the mass (kg) of the tag and a is the acceleration (*g*).

Statistical analyses

Linear mixed-effects models were conducted in R Studio V 1.2.1135 within the ‘Lme4’ package in order to investigate how the period between acceleration peaks, gait and body mass influenced peak accelerations across species. Additionally, we investigated how travel speed (covariate), body mass (covariate), collar mass as a percentage of carrier body mass (fixed factor) and gait (fixed factor) influence peak accelerations and consequent forces exerted by the tags. The influence of dog body mass and collar mass as a percentage of carrier body mass on gait-specific travel speed and the period between peak accelerations was also investigated. Dog ID was included as a random factor in all models. All potential interaction effects were first investigated and a step-wise back-deletion of non-significant interaction terms was conducted. Standard model diagnostics were conducted in order to ensure that model assumptions were met (examining q-q plots and plotting the residuals against fitted values) and data transformations were conducted in order to meet assumptions where appropriate. Outputs of the final models are reported. The F statistic and marginal and conditional R^2^ were determined using the ‘car (3.0-5)’ and ‘MuMIn (1.46.6)’ packages, respectively. Coefficients for best-fit lines in the figures were extracted from the outputs of the models
