## Supplementary table 1 for "Forces experienced by instrumented animals depend on lifestyle"

Table S1.

Percentage time the tags were exposed to accelerations (vector sum) greater than the values specified in brackets according to species and gait. Note that pine martens only moved substantially by bounding.

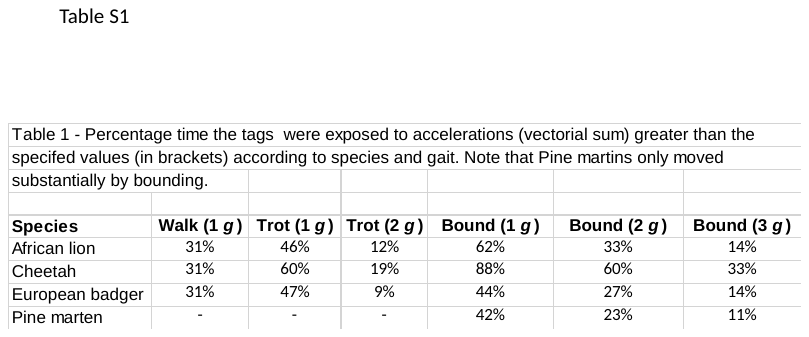
