## Supplementary table 2 for "Forces experienced by instrumented animals depend on lifestyle"

Table S2.

Mean peak accelerations (SD) [vectorial sum – *g*] per stride measured using a collar-attached tag for four wild animal species as a function of activity. Note that pine martins only moved substantially by bounding.

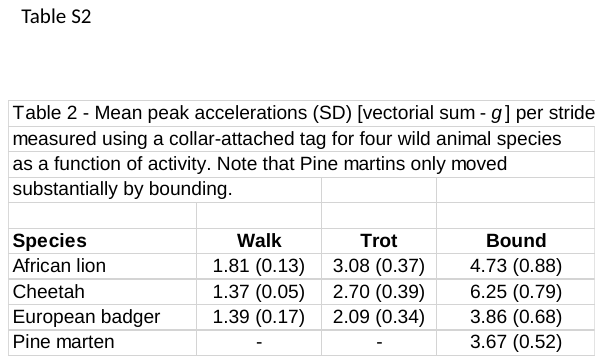
