## Supplementary table 3 for "Forces experienced by instrumented animals depend on lifestyle"

Table S3.

Final outputs of linear mixed-effects models conducted to investigate the factors influencing peak accelerations in wild carnivores. Data were logged to ensure that model residuals were normally distributed.

| **Parameter** | **Final model terms** | ***F*** | **DF** | ***P*** | ***R^2^ fixed*** | ***R^2^total*** |
| --- | --- | --- | --- | --- | --- | --- |
| log peak acceleration | log(period)  gait  body mass | 0.013  1083.07  3.00 | 1, 210.40  2, 208.39  1, 18.79 | 0.908  <0.0001  0.100 | 0.88 | 0.92 |
| period | body mass  gait  bodymass x gait | 99.98  114.50  76.27 | 1, 14.46  2, 210.03  2, 209.29 | <0.0001  <0.0001  <0.0001 | 0.73 | 0.76 |
