## Supplementary table 4 for "Forces experienced by instrumented animals depend on lifestyle"

**Table S4.**

Summary of the outputs of linear mixed-effects models conducted to investigate how dog body mass (*M_b_*), collar mass as a percentage of carrier mass (%*M_b_*), travel speed (*U*) and gait influence different parameters. Dog ID was included as a random effect in all models. Final models following the removal of non-significant interaction terms are reported

| **Model** | **Parameter** | **Final model terms** | ***F*** | **DF** | ***P*** | ***R^2^_f_*** | ***R^2^*_t_** |
| --- | --- | --- | --- | --- | --- | --- | --- |
| A | peak acceleration  (*g*) | period  gait  body mass  period x gait | 10.81  340.88  0.21  7.92 | 1, 496.55  2, 509.97  1, 10.15  514.49 | 0.001  <0.0001  0.658  <0.0001 | 0.19 | 0.54 |
| B | period between peaks (s) | body mass  gait  body mass x gait | 3.37  47.54  9.90 | 1, 9.98  2, 508.02  2, 508.04 | 0.096  <0.0001  <0.0001 | 0.20 | 0.54 |
| C | speed (m s^-1^) | body mass  % body mass  gait  body mass x gait | 5.64  1.67  1500.6140.15 | 1, 9.96  3, 507.49  2, 505.04  2, 505.10 | 0.039  0.172  <0.0001  <0.0001 | 0.82 | 0.86 |
| D | peak acceleration (*g*) | speed  % body mass  gait  body mass  speed x % body mass  % body mass x gait | 40.59  35.15  21.15  3.51  4.44  4.33 | 1, 506.91  3, 499.60  2, 502.31  1, 10.12  3, 500.77  6, 498.57 | <0.0001  <0.0001  <0.0001  0.090  0.004  0.0002 | 0.66 | 0.74 |
| E | force as % body mass | body mass  % body mass  gait  body mass x % body mass  body mass x gait  % body mass x gait | 4.62  126.44  75.28  37.68  7.38  16.10 | 1, 9.98  2, 356.97  2, 356.01  2, 356.90  2, 356.03  4, 356.01 | 0.057  <0.0001  <0.0001  <0.0001  <0.0001  <0.0001 | 0.38 | 0.84 |
