## Supplementary figure legends for "Forces experienced by instrumented animals depend on lifestyle"

Fig. S1.

Peak amplitudes of (the vectorial sum of) accelerations versus period between peaks for walking (orange), trotting (blue) and bounding (pink) in twelve domestic dogs ranging 2-45 kg. Dat

Fig. S2.

Peak amplitudes of (the vectorial sum of) accelerations versus body mass for twelve dogs ranging 2-45 kg in body mass. A) walking. B) trotting. C) bounding. Data points include the four greatest peaks in acceleration per 20 m trial per dog and are coloured according to tag mass as a percentage of carrier body mass; 0% (grey), 1% (yellow), 2% (light blue) and 3% (dark blue).

**Fig. S3.**

Forces exerted by the tags as a percentage of dog body mass for twelve dogs ranging 2-45 kg in body mass. Data points show walking (triangles), trotting (circles) and bounding (squares), and are coloured according to tag mass as a percentage of carrier body mass; 1% (yellow), 2% (light blue) and 3% (dark blue). The dashed grey line indicates the relative force of a tag of 3% body mass at 1 *g.*
