## Supplementary figures and images for "Forces experienced by instrumented animals depend on lifestyle"

### Supplementary fig 2

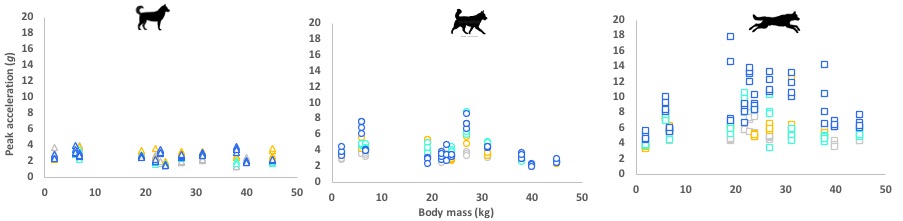

### supplementary fig 3

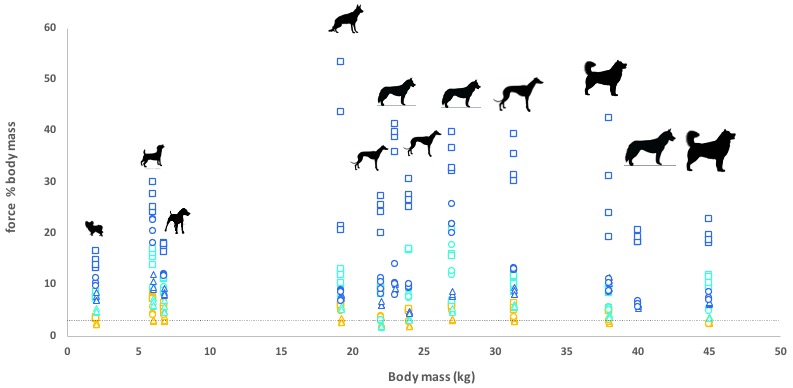

### Supplementary figure 1

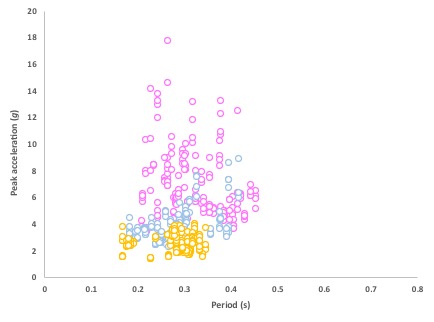
